# *Trithorax* balances ISC fate decisions via Ptx1-mediated repression of EE specification

**DOI:** 10.1101/2025.11.21.689716

**Authors:** Virginia Fasoulaki, Aristides G. Eliopoulos, Alexandros Galaras, Vasiliki Theodorou, Pantelis Hatzis, Christos Delidakis, Zoe Veneti

## Abstract

Adult stem cells must coordinate transcriptional programs with external cues to maintain tissue homeostasis. In the *Drosophila* midgut, intestinal stem cells (ISCs) generate enterocytes (ECs) or enteroendocrine (EE) cells through Notch-dependent fate decisions, but how chromatin regulators influence this balance remains unclear. In this study we identified the Trithorax gene (*trx*) as a key factor that safeguards ISC lineage fidelity. Trx depletion biases ISCs toward EE differentiation without affecting proliferation, a phenotype exacerbated during aging or DSS-induced damage. Transcriptomic and chromatin profiling revealed the homeobox transcription factor *Ptx1* as a Trx-dependent target. *Ptx1* knockdown phenocopies Trx loss, whereas *Ptx1* overexpression reverts *trx*-RNAi-induced EE overproduction, establishing *Ptx1* as a critical mediator of Trx function. These findings support a model in which Trx constrains the *scute–prospero* axis through *Ptx1*-mediated repression, thereby limiting inappropriate EE specification and maintaining ISC plasticity.

## Introduction

Precise control of stem cell fate decisions is essential for tissue homeostasis and regeneration. In the adult *Drosophila* midgut, intestinal stem cells (ISCs) continually replenish the epithelium by generating absorptive enterocytes (ECs) and secretory enteroendocrine (EE) cells through transient progenitor cells called enteroblasts (EBs) and enteroendocrine progenitors (EEPs) respectively, a process coordinated by canonical signaling pathways and chromatin-based regulatory mechanisms. ISC decisions are primarily dictated by Notch signaling, which promotes EC differentiation, and by the proneural factor Scute, which drives expression of the alternative EE cell fate via expression of the EE determinant Prospero (Pros) (Ohlstein and Spradling, 2007; Zeng and Hou, 2015; Chen et al., 2018). Together with daughterless (Da), which antagonizes both differentiation trajectories and promotes ISC self-renewal (Puig-Barbe et al., 2025), these pathways provide a general framework for ISC lineage specification, yet how epigenetic regulators interface with them to ensure balanced fate choices is only beginning to emerge.

Chromatin regulators of the Polycomb group (PcG) and Trithorax group (TrxG) are central antagonistic players that maintain cellular identity during development and in adult tissues. PcG proteins mediate transcriptional repression, primarily through trimethylation of histone H3 lysine 27 (H3K27me3) catalyzed by the PRC2 subunit Enhancer of zeste [E(z)], and through PRC1-mediated chromatin compaction and ubiquitination of histone H2A at Lys119 (H2AK119ub1). In contrast, TrxG proteins counteract PcG repression by sustaining transcriptional activity at target loci, via histone H3 lysine 4 (H3K4) methylation and chromatin remodeling. The balance between these opposing activities establishes stable yet flexible gene expression states, enabling stem cells to both preserve identity and respond to differentiation cues (Kuroda et al., 2020; Brand et al., 2019; Schuettengruber et al., 2017).

Recent studies highlighted roles for both groups of epigenetic regulators in Drosophila ISC regulation. TrxG member Trithorax-related (Trr) cooperates with the nucleosome remodeler Kismet to restrain ISC proliferation by activating the EGFR antagonist *Cbl* (Gervais et al., 2019). PcG proteins also play pivotal roles: the PRC1 subunit Polycomb (Pc) ensures proper fate outcomes, with increased Pc activity in aging midguts biasing differentiation toward the EE lineage (Tauc et al., 2021). Similarly, E(z) represses EC-promoting factors, such as *miR-8,* to maintain ISC identity (Veneti et al., 2024), and PRC1 components Psc, Su(z)2, Sce, and Ph protect ISCs from turning tumorous by regulating JAK/STAT and Toll signaling (Joly et al., 2025). Given the evolutionary conservation of PcG and TrxG proteins, and their frequent deregulation in cancer, including colorectal cancer, defining their precise functions in *Drosophila* ISCs will provide critical insights into how chromatin-based mechanisms safeguard epithelial stem cell identity and prevent malignant transformation.

In this study, we uncover a critical role for the *trithorax* gene (*trx*), in constraining ISC lineage decisions. Using lineage-specific knockdown and clonal analysis, we show that *trx* loss leads to excessive EE differentiation without affecting ISC proliferation. Mechanistically, we identify a set of Trx-activated transcription factors—most notably the paired-like homeobox gene *Ptx1*—that function to repress the pro-EE Achaete-Scute Complex (AS-C)–Pros axis. These results highlight a previously unrecognized chromatin-based mechanism that constrains lineage decisions in a regenerating epithelium and underscore the coordinated interplay between chromatin regulators, transcription factor codes, and signaling pathways in controlling adult stem cell behavior.

## Results

### Trithorax knockdown promotes enteroendocrine differentiation

To assess the role of Trithorax (trx) in adult intestinal homeostasis, we specifically knocked down trx in intestinal stem cells (ISCs) and progenitor cells, including enteroblasts (EBs) and enteroendocrine precursors (EEPs), using the esg-GAL4; tub-GAL80^ts^; UAS-GFP system (Micchelli and Perrimon, 2006) with three different trx RNAi lines. Cell types were distinguished by immunofluorescence staining for Delta (Dl) (ISCs), Prospero (Pros) (EEPs/EEs), GFP (esg-positive cells), and DAPI (nuclei).

Knockdown with all three RNAi lines for 7 days resulted in a significant increase in Pros⁺ cells, indicative of enhanced EE differentiation, particularly in the posterior midgut (**Fig. 1A–B, 1G, Suppl. Fig. 1A–D**). In contrast, Dl⁺ ISC and GFP⁺ progenitor numbers remained unchanged, indicating that the EE increase was not due to expansion of the progenitor pool (**Fig. 1H–I**).

**Figure 1.**
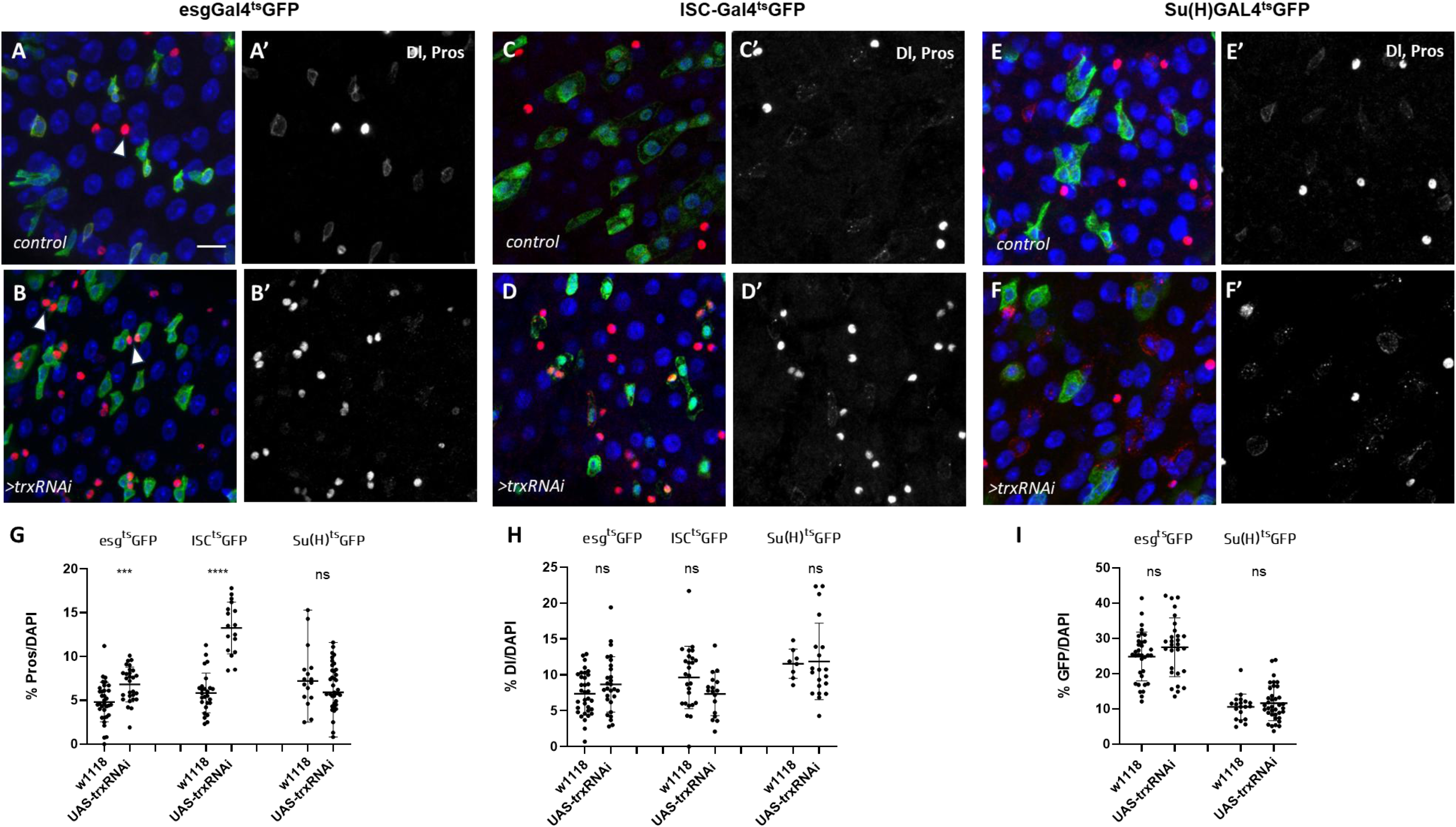
Loss of trx in ISCs skews cell divisions toward the EE fate Flies carrying cell-specific Gal4 drivers were crossed either to UAS-trx-RNAi to deplete trx or to w1118 to generate the corresponding control genotype. Young mated female flies were dissected 7 days post temperature shift at 29°C and immunostainings were performed for the ISC marker Delta (Dl, cytoplasmic puncta) and the enteroendocrine (EE) cell marker Prospero (Pros, solid nuclear staining). Scale bar: 20 µm. (A, B) Immunostaining of posterior midguts from control and UAS-trxRNAi (BDSC#33703) flies carrying the ISC/EB/EEP driver esg^ts^ > GFP. Number of midguts: n = 12 (control) and n = 12 (trxRNAi). Arrows mark enteroendocrine cells (EE, Pros⁺). (C, D) Immunostaining of UAS-trxRNAi expression (BDSC#33703) using the ISC-specific esg-Gal4^ts^ >GFP; Su(H)GBE-Gal80 driver. Number of midguts: n = 10 (control) and n = 8 (trxRNAi). (E, F) Immunostaining of RNAi-mediated knockdown of trx (v37715) using the EB-specific Su(H)GBE-Gal4^ts^ driver. Number of midguts: n = 8 (control) and n = 12 (trxRNAi). (G) Quantification of Pros+ cells using the different Gal4 drivers expressed as percentage of total DAPI-stained cells. For each genotype, two to four zoom-in images from each midgut were randomly selected and are represented as a single dot in the graphs. Statistical analysis was performed using a two-tailed unpaired t-test. Data are presented as mean ± standard deviation (SD). (H) Quantification of Dl-positive intestinal stem cells across the Gal4 drivers crossed with w1118 or UAS-trxRNAi (BDSC#33703) shown in the images, expressed as the percentage of total DAPI-stained cells. Each dot represents one zoom-in image. Statistical significance was determined using a two-tailed unpaired t-test. Data are shown as mean ± SD. (I) GFP⁺ cells were quantified in control and trxRNAi midguts using esg^ts^Gal4 (marking all progenitor cells) and Su(H)^ts^Gal4 (marking enteroblasts), expressed as a percentage of total DAPI⁺ cells. Each dot corresponds to one zoom-in image. Statistical significance was assessed using a two-tailed unpaired t-test. Data are presented as mean ± SD. For trx knockdown, BDSC#33703 was used for the esg^ts^Gal4 crosses, whereas v37715 was used for the Su(H) ^ts^Gal4 crosses.

To define the cellular origin of this phenotype, we used additional GAL4 drivers to knockdown trx selectively in ISCs or EBs (Wang et al., 2014; Zeng et al., 2010). ISC-specific trx depletion recapitulated the EE increase without affecting ISC numbers (**Fig. 1C–D, 1G-H,**), whereas EB-specific knockdown had no effect on ISCs, EE or EB numbers (**Fig. 1E–F, 1G-I**). These results indicate that trx acts cell-autonomously in ISCs to prevent excessive EE commitment rather than misrouting EBs to the alternative EE fate.

### Trx loss does not affect ISC proliferation

To investigate whether trx regulates ISC proliferation, we first quantified mitotic activity in control and esg^ts^>trx RNAi midguts using phospho-Histone H3 (pH3) staining. A notable increase in pH3⁺ cells was observed in Trx-depleted intestines (**Fig. 2A–C**). However, marker analyses indicated that many of these dividing cells were EEPs rather than ISCs (**Suppl. Fig. 2A-A’**), suggesting that the bulk increase in proliferation may reflect the EE lineage bias induced by trx knockdown, as EEPs undergo a terminal division shortly after they form, in contrast to non-dividing EBs.

**Figure 2.**
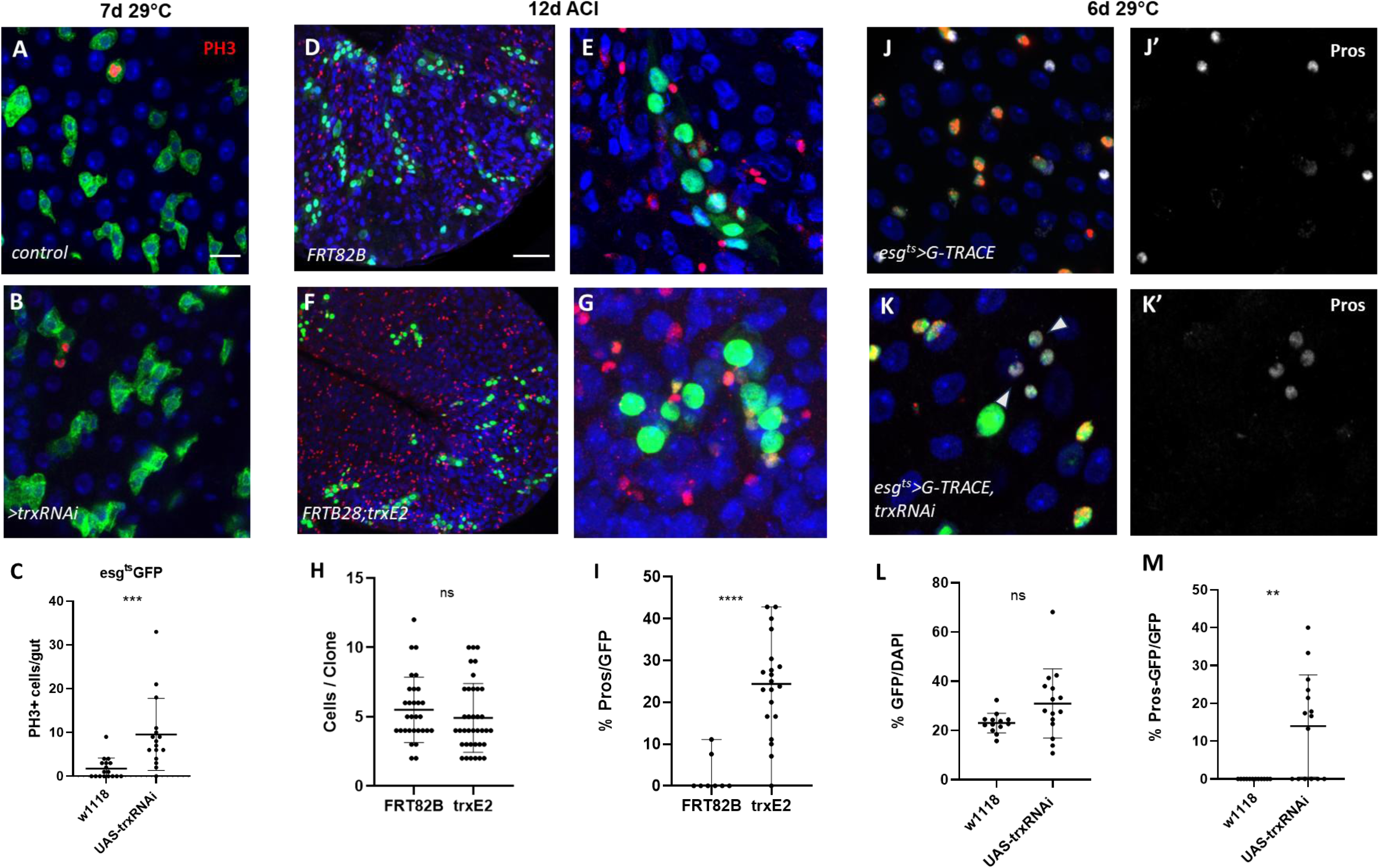
Trx loss does not affect ISC proliferation (A, B) Immunostaining of posterior midguts from control (w1118) and UAS-trxRNAi flies (BDSC#33703) using the ISC/EB/EEP driver esg^ts^GFP 7 days after temperature shift to 29°C. Mitotic cells were labeled with phospho-Histone H3 (PH3,nuclear red). Scale bars: 20 µm. (C) Quantification of pH3+ positive cells representing mitotic cells in control and trx-RNAi midguts. Cells were counted from whole midguts and each dot represents a whole midgut in the quantification graph. n = 17 for control flies and n = 16 for trx-RNAi flies. Data were pooled from three individual experiments. Data represented as mean ± SD. Statistical significance was assessed using a two-tailed unpaired Student’s t-test. (D–G) MARCM clonal analysis comparing control (D) and trx deficient (F) clones 12 days after heat-shock induction. Immunostaining was performed for the markers Delta (Dl) and Prospero (Pros) to label ISCs and EE cells, respectively. Clones were induced by heat shock at 37 °C for 1 h. Flies were then maintained at 25 °C throughout the clonal expansion period. Higher-magnification views from control and trx mutant clones are shown in (E) and (G), respectively. Scale bars: 50 µm (overview D, F), 20 µm (zoom-in E, G). (H) Quantification of GFP+ cells per clone, showing clonal growth in control and trx mutant clones. A two-tailed unpaired t-test was used to evaluate statistical significance. Values are shown as mean ± SD. (I) Quantification of Pros+ cells expressed as a percentage within GFP-marked clones to assess the EE numbers produced upon trx loss. Statistical analysis was conducted with a two-tailed unpaired t-test. Data represented as mean ± SD. (J-K) Representative immunofluorescence images of midguts from control (J-J’) and UAS-trxRNAi (K-K’) flies (BDSC#33703) using the G-Trace system 6 days after temperature shift to 29°C. Progenitor cells are marked by both RFP and nuclear EGFP. Newly formed ECs are identified as large polyploid EGFP⁺ cells and newly formed EEs as Pros⁺, EGFP⁺ cells. DAPI marks nuclei. Scale bars: 20 µm. (L) Newly produced cells marked by nuclear GFP were quantified and expressed as a percentage of total DAPI-stained cells 6 days after the temperature shift to 29 °C to assess ISC proliferation in control and trxRNAi midguts. Data represent mean ± SD from n = 5 midguts per genotype. Statistical significance was determined using unpaired two-tailed Student’s t-test. (M) Quantification of newly formed enteroendocrine cells (EEs), shown as the percentage of Pros⁺GFP⁺ cells relative to the total number of GFP⁺ cells in control and trx-RNAi midguts. Data represent mean ± SD from n = 5 midguts per genotype. Statistical significance was determined using unpaired two-tailed Student’s t-test.

To further assess proliferative capacity and lineage behavior, we generated trx mutant clones using MARCM (Wu and Luo, 2006). GFP-marked homozygous trx clones were induced on the FRT82B chromosome arm using the amorphic allele trx^E2^ in an otherwise heterozygous background. Clone size and Dl^+^ cells, were comparable between trx mutant and control clones (**Fig. 2D–H, Suppl. Fig. 2B**), indicating that Trx is dispensable for ISC self-renewal and expansion under homeostatic conditions. However, trx mutant clones more frequently contained multiple Pros⁺ cells (**Fig. 2I**), confirming a lineage bias toward EE fate in the absence of Trx.

We also performed G-TRACE lineage analysis (Evans et al., 2009) in esg^ts^>trx RNAi midguts. In this system, esg-driven GAL4 activated RFP in ISCs and progenitor cells and permanently labeled their progeny with nuclear GFP following UAS-FLP-mediated STOP cassette excision. Six days post-induction, we quantified all cell types and observed no increase in total GFP⁺ lineage size (**Fig. 2J-L**), but a clear elevation in the proportion of newly-born Pros⁺ cells (**Fig. 2M**), reinforcing that Trx loss alters cell fate without impacting proliferation.

Collectively, these findings demonstrate that Trx acts to restrict EE differentiation without affecting ISC proliferation, highlighting its cell-autonomous role in lineage specification.

### Trithorax restricts EE differentiation under stress conditions

As intestinal injury and aging perturb intestinal proliferation and differentiation, we asked whether Trx also modulates EE fate in these contexts. In mammals, oral dextran sulfate sodium (DSS) physically compromises the colonic mucosal barrier and induces ulcerative-colitis–like pathology (Gkouskou et al., 2016). In *Drosophila*, DSS feeding greatly increases the numbers of GFP^+^ progenitors and pH3^+^ mitotic cells relative to controls, consistent with damage-induced regenerative responses, but does not affect Pros⁺ EE cells (Amcheslavsky et al., 2009). After 48 hours of DSS feeding, trx-depleted midguts exhibited a slight increase in pH3⁺ cells (**Fig. 3A-C**) but a considerably stronger augmentation in Pros⁺ cells (**Fig. 3D–F**), indicating that Trx restricts excessive EE differentiation during regenerative stress, as it does in homeostatic conditions. Lineage tracing after 5 days of RNAi induction followed by 24h of DSS feeding confirmed the aforementioned bias of ISC fate decisions in esg^ts > trx RNAi midguts (**Fig. 3G-J**).

**Figure 3.**
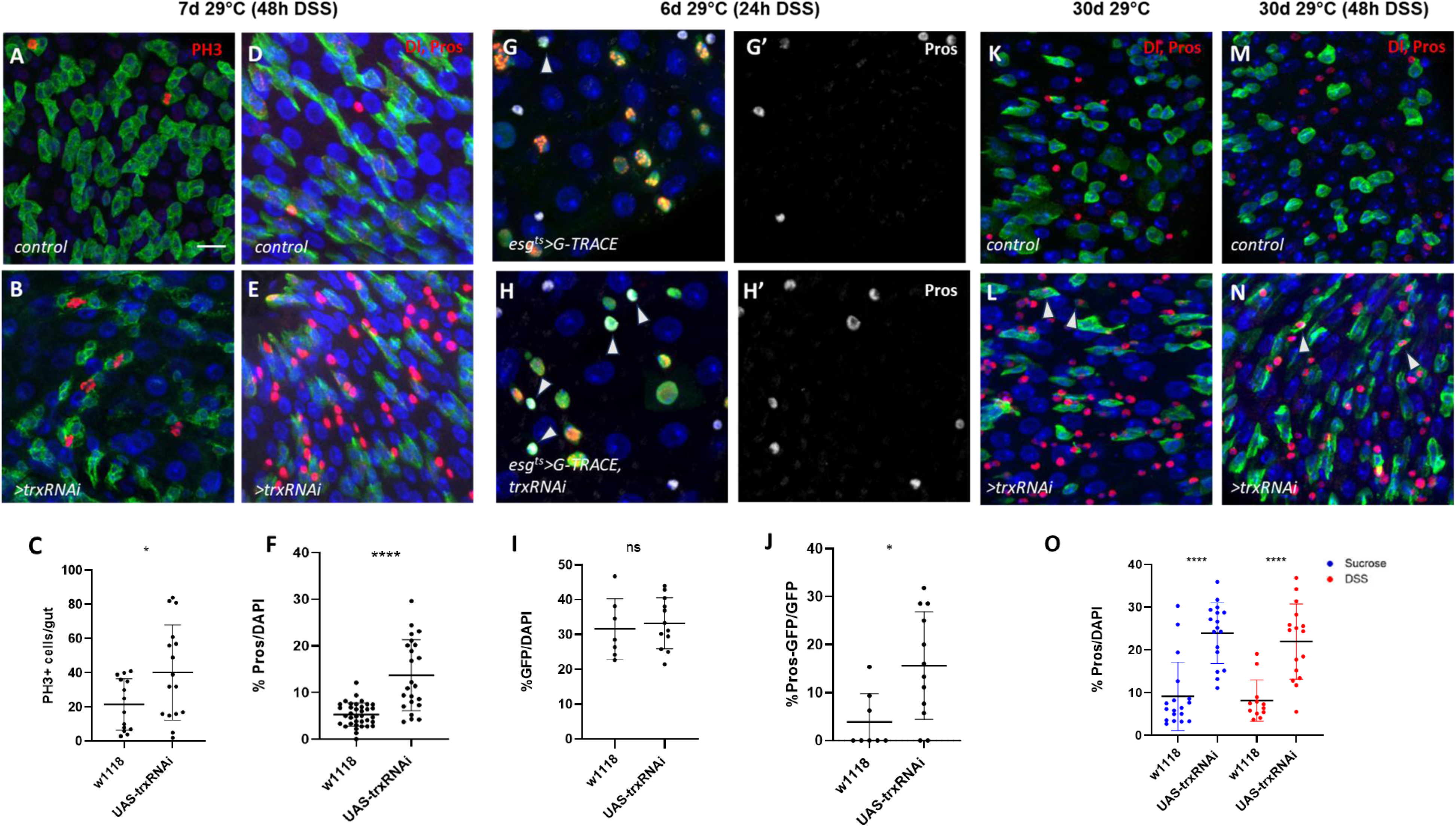
Trx loss drives excess EE differentiation in stress and aging (A-C) Representative confocal images of posterior midguts from control (A) and UAS-trxRNAi (B) flies (BDSC#33703), using the esg-Gal4tsUAS-GFP system. Flies were maintained at 29 °C for a total of 7 days to induce RNAi expression, and were fed DSS during the final 48 hours to elicit an acute inflammatory response. Midguts were stained and quantified for phosphorylated histone H3 (PH3, nuclear red) to assess mitotic activity. Counts were obtained from entire midguts. Each dot represents a whole midgut measurement; Data were pooled from three independent experiments and correspond to n=14 for control and n=16 midguts for trxRNAi. Data are shown as mean ± SD. Statistical analysis was performed using a two-tailed unpaired t-test. Scale bars: 20 µm. (D-F) Midguts from control (D) and trxRNAi (E) flies (BDSC#33703) using the esg-Gal4 driver were stained for Delta (Dl, cytoplasmic puncta) and Prospero (Pros, red nuclear) to evaluate the numbers of intestinal stem cells and enteroendocrine cells, respectively. Flies were maintained at 29 °C for a total of 7 days, with DSS feeding during the final 48 hours to model an acute inflammatory condition. Quantification of Pros+ cells (F) is expressed as a percentage of total DAPI-stained cells. Statistical significance was performed using a two-tailed unpaired t-test. Data are presented as mean ± standard deviation (SD). Scale bars: 20 µm. (G-H) Representative immunofluorescence images of midguts from control (G–G’) and trx-RNAi (H–H’) flies (BDSC#33703) analyzed with the G-TRACE system after 6 days of RNAi induction at 29°C with DSS feeding during the final 24 hours to trigger ISC divisions. Progenitor cells are labeled by RFP and nuclear EGFP, while newly differentiated ECs appear as large polyploid EGFP⁺ cells and newly differentiated EEs as double Pros⁺ EGFP⁺ cells. Nuclei are visualized with DAPI. Scale bars: 20 µm. (I-J) Quantification of ISC proliferation (I) and EE differentiation (J) in control and trx-RNAi midguts after 6 days at 29 °C, with DSS administered during the final 24h. Proliferation was evaluated by measuring newly produced cells marked by nuclear GFP, expressed as a percentage of total DAPI-stained cells (I), while EE differentiation was assessed as the proportion of newly formed EEs (Pros⁺GFP⁺) relative to the total GFP⁺ population (J). Data represent mean ± SD; statistical significance was determined using an unpaired two-tailed Student’s t-test. (K-L) Confocal images of posterior midguts from aged control and trx-RNAi flies (BDSC#33703) maintained at 29 °C for 30 days, with flies transferred to fresh food every 3–4 days. Nuclei were stained with DAPI (blue). Pros was used to label enteroendocrine (EE) cells and recently produced EEs were identified as Pros⁺GFP⁺ (EEPs, indicated by arrows). Data are shown as mean ± SD. Scale bars: 20 µm. (M-N) Posterior midgut immunostainings from aged control and trxRNAi flies (BDSC#33703) subjected to DSS-induced epithelial damage. Flies were kept at 29 °C for 30 days and transferred to fresh food every 3–4 days. On day 28, flies were fed DSS for 48 h to stimulate ISC divisions in the aged epithelium. Nuclei were visualized with DAPI (blue). EE cells were marked with Pros, and newly generated EEs (EE progenitors) were identified as Pros⁺GFP⁺ cells (arrows). Data are presented as mean ± SD. Scale bars: 20 µm. (O) Percentage of Pros⁺ cells relative to total DAPI-stained nuclei in the aged midgut epithelium of control and trxRNAi flies under normal feeding (30d 29 °C) or following a 2-day DSS treatment (30d 29 °C + DSS). Data are mean ± SD and statistical significance was determined by unpaired two-tailed Student’s t-test.

We next analyzed aged animals, which naturally accumulate misdifferentiated cells, particularly EEs, and display aberrant ISC activity (Tauc et al., 2021). In 30-day-old midguts, Pros⁺ cells were detectably elevated relative to young flies, consistent with age-associated EE overproduction. Strikingly, trx knockdown further exacerbated EE accumulation (**Fig. 3K–L, 3O**), including Pros⁺GFP⁺ double-positive EEPs (arrows), indicating sustained production of EEs. In contrast, DSS treatment of aged *trx*-depleted midguts did not further increase the proportion of Pros⁺ cells (**Fig. 3M–O**), suggesting that aged intestines, which are already hyperplastic and inflamed, are less responsive to additional DSS-induced EE differentiation, although newly formed transient EEPs could still be detected (**Fig. 3N**, arrows and **Suppl. Fig. 2C**).

Together, these findings demonstrate that Trx restrains EE lineage commitment not only during homeostasis but also under stress conditions such as inflammation and aging, when fate regulation is particularly susceptible to disruption.

### Trithorax functions upstream of proneural /Notch network

To investigate the mechanism by which Trx limits EE specification, we first confirmed efficient trx knockdown in FACS-sorted esg⁺ progenitors using qRT-PCR. Trx transcript levels were reduced in esg^ts^>trx RNAi midguts by >60% compared to controls (**Fig. 4A**).

**Figure 4.**
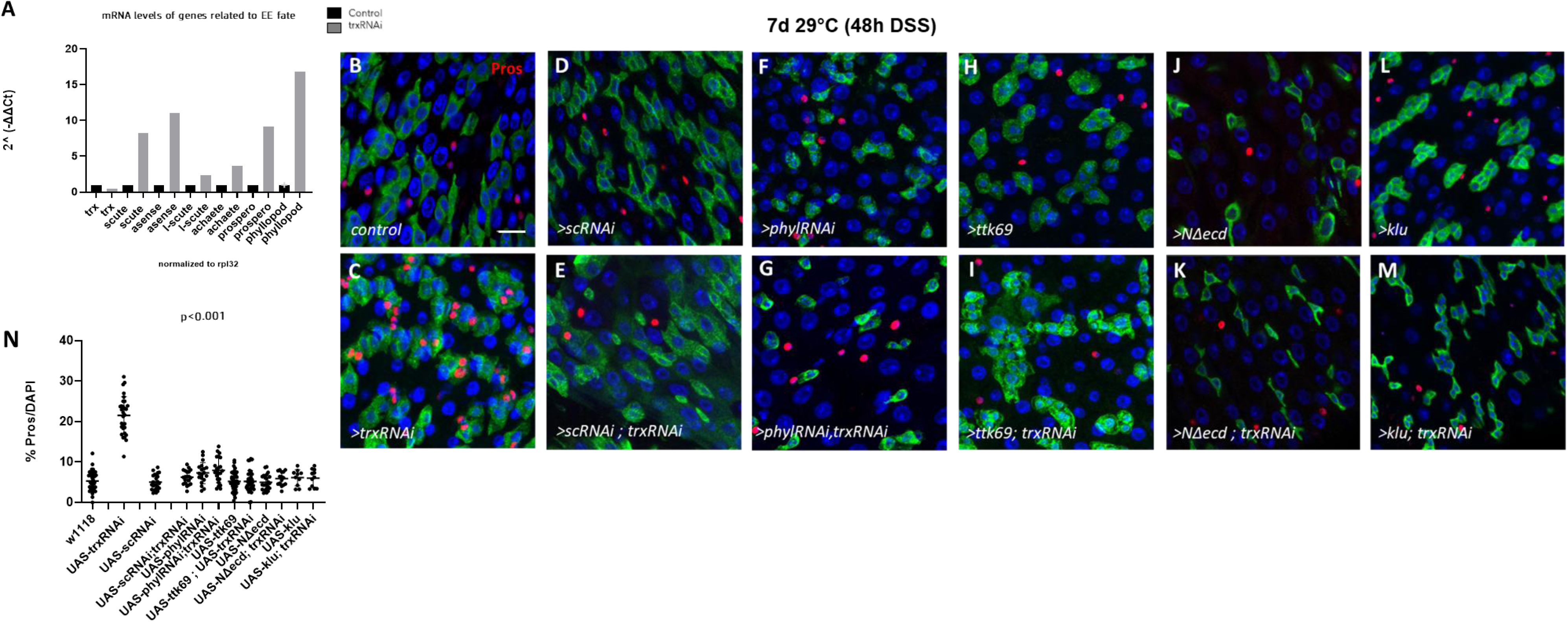
Trithorax functions upstream of proneural /Notch network (A) RT-qPCR analysis of mRNA expression levels of EE-related genes (scute, asense, phyllopod, prospero) in progenitor cells depleted of trx compared to controls. Virgin female flies carrying the esg-Gal4tsGFP driver were crossed to UAS-trx-RNAi (BDSC#33703) to induce trx knockdown, or to w1118 to generate the corresponding control genotype. Young mated females were dissected 7 days after the temperature shift to 29°C. Total RNA was extracted from two biological replicates of FACS-sorted GFP⁺ progenitors, isolated from 120 control and 120 trx-RNAi midguts per replicate. Expression levels were normalized to rpl32, as indicated. (B-M) Representative images of posterior midguts from young female flies expressing the indicated RNAi constructs under the control of *esg*-Gal4^ts^GFP driver, following 7 days of total induction at 29 °C and 48 hours of DSS feeding to trigger ISC divisions. BDSC#33703 was used as the RNAi line to knock down *trx*. Flies carrying only the single-gene manipulations were obtained from the same parental crosses as those used to produce the combinatorial trxRNAi genotypes, ensuring a consistent genetic background. All genotypes were raised and handled under identical environmental conditions and DSS was administered uniformly prior to dissection. Enteroendocrine cells (EEs) were visualized by immunostaining for the EE marker Prospero (Pros, red) and nuclei were counterstained with DAPI (blue). Scale bar; 20 μm, applies to all images. (N) Quantification of EE content expressed as the percentage of Pros+ cells over total DAPI+ nuclei. Data represent mean ± SD. All experimental genotypes showed statistically significant differences compared to control (p < 0.001, two-tailed unpaired t-test).

We next examined the expression of genes that drive EE differentiation (Yin and Xi, 2018; Chen et al.,2018). qRT–PCR revealed upregulation of the proneural factor scute (sc) and its downstream effectors prospero (pros) and phyllopod (phyl) in Trx-depleted progenitors (**Fig. 4A**), suggesting that Trx normally represses this EE-promoting transcriptional program.

To test whether these factors are functionally required for the Trx RNAi phenotype, we performed epistasis experiments. Simultaneous knockdown of scute in the trx RNAi background abolished EE overproduction (**Fig. 4B–E, N**), demonstrating that scute is essential for the EE bias induced by Trx loss. Similar results were obtained for additional genes implicated in *scute* regulation. The transcriptional repressor Ttk69 (Wang et al., 2015) normally inhibits *scute* expression, while Phyllopod (Phyl) targets Ttk69 for degradation, relieving repression and reinforcing *scute* activation through a self-amplifying feedback loop (Yin and Xi, 2018). We found that phyl RNAi or ttk69 overexpression were epistatic to trx knockdown (**Fig. 4F–I, N**).

We next asked whether promoting the alternative EB–EC fate could counteract the EE bias. Overexpression of an active form of Notch promotes ISC differentiation to EC (Ohlstein and Spradling, 2007) When we expressed UAS-NΔecd together with trx-RNAi, EE overproduction was strongly suppressed (**Fig. 4J–K, N**). Consistent with this, overexpression of the Notch effector klumpfuss (klu) (Korzelius et al., 2019) also rescued the phenotype (**Fig. 4L–N**), indicating that forced EB–EC specification can override the loss of Trx function.

Together, these findings place Trx upstream of both EE-promoting proneural factor scute and the Notch signaling network. By repressing the AS-C–Pros axis, Trx restricts inappropriate EE fate acquisition and thereby ensures that most ISC divisions adopt the EB-EC trajectory.

### Chromatin and transcriptome profiling of putative Trx-regulated genes

To identify genes regulated by Trx that may mediate its role in suppressing EE differentiation, we performed RNA-seq and ATAC-seq on FACS-sorted esg⁺ stem / progenitor cells from control and trx RNAi adult midguts (7d exposure), under both homeostatic (sucrose-fed) and regenerative (DSS-treated) conditions (**Fig. 5A**).

**Figure 5.**
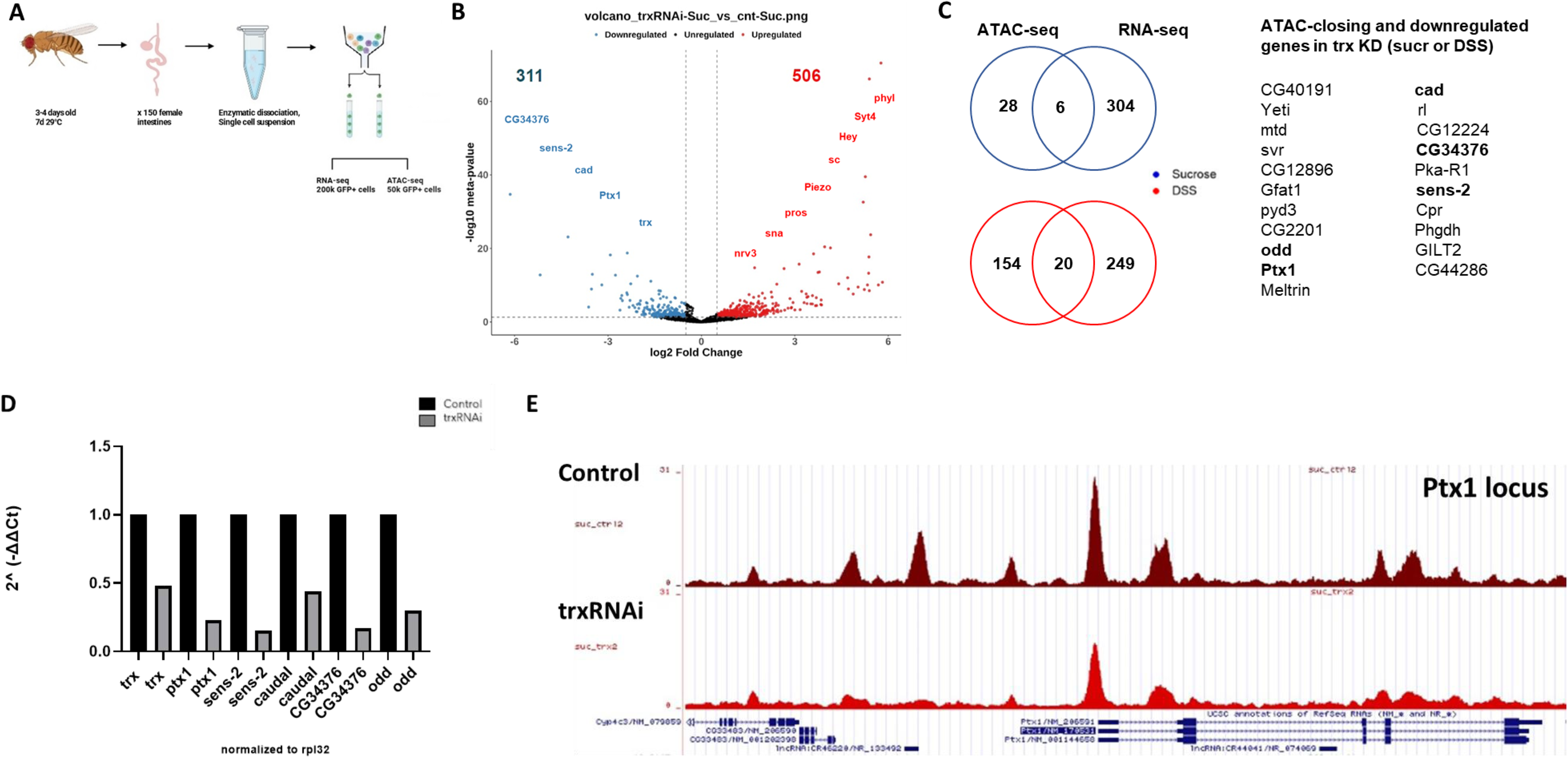
Chromatin and transcriptome profiling of putative Trx-regulated Genes (A) Experimental workflow for midgut dissection and FACS-based isolation of GFP⁺ ISC progenitors from control and trxRNAi midguts. Flies were generated by crossing esg-Gal4, tub-Gal80ts, UAS-GFP (esgtsGFP) to w1118 to obtain controls, or to UAS-trxRNAi (BDSC#33703) to induce trx knockdown in intestinal progenitor cells. Young mated females were dissected 7 days after the temperature shift to 29 °C. The same workflow was used for both homeostatic and DSS-treated conditions; for DSS experiments, flies were maintained at 29 °C for a total of 7 days, with DSS administered during the final 48 h. For each biological replicate, GFP⁺ cells from ∼150 dissected flies per genotype were isolated using established sorting procedures, and cells from the same replicate were used for both ATAC-seq and RNA-seq. Two biological replicates were used for ATAC-seq, and three biological replicates were used for RNA-seq. (B) Volcano plot of RNA-seq data showing significantly differentially expressed genes upon trx knockdown in ISC progenitor cells under homeostatic conditions. Significantly up- and down-regulated transcripts are highlighted in red and blue, respectively. Red labels correspond to selected up-regulated genes associated with EE fate, while blue labels denote trx (confirming its efficient knockdown) together with down-regulated transcription factors. (C) Venn diagrams illustrating the overlap between downregulated genes (RNA-seq) and genes with reduced chromatin accessibility (ATAC-seq) in trxRNAi progenitors under sucrose feeding (top, red) and DSS treatment (bottom, blue). The gene list on the right summarizes loci showing both decreased RNA expression and reduced chromatin accessibility in trxRNAi progenitors under either condition. Transcription factors within this set are highlighted in bold. (D) mRNA expression analysis of transcription factor genes tested as candidate repressors of EE fate. mRNA levels were analyzed by RT-qPCR in progenitor cells depleted of trx compared with controls. Flies carrying the *esg*-Gal4^ts^GFP driver were crossed to UAS-trx-RNAi (*BDSC#33703)* or to w1118 to generate the corresponding control. Total RNA was extracted from one biological replicate of FACS-sorted GFP+ progenitors, each consisting of cells from 130 control or 130 trx-RNAi midguts, and expression levels were normalized to rpl32. (E) UCSC Genome Browser view of the Ptx1 locus showing normalized ATAC-seq signal in control and trxRNAi ISC progenitors under sucrose feeding. trx knockdown leads to an overall reduction in chromatin accessibility across the Ptx1 locus, extending into upstream regulatory regions.

While Trx is canonically known as a transcriptional activator via H3K4 methylation (Tie et al., 2014; Rickels et al., 2016), our analysis revealed that many genes were upregulated and displayed increased chromatin accessibility in trx-depleted progenitors (**Fig 5B, Suppl. Fig. 3A-B**). This likely reflects indirect effects, such as the derepression of the EE-lineage program following Trx loss. Based on the results presented so far, we hypothesized that a putative repressor of scute (and/or other pro-enteroendocrine regulators) is kept in an active state via Trx-mediated chromatin modification. Therefore, we focused on the subset of genes that were both downregulated and showed reduced chromatin accessibility upon trx depletion (**Fig. 5C**).

We used a cutoff of log_2_FC<-0.5 and FDR<0.05 for both RNA expression and ATAC data. Applying these criteria in both homeostatic and DSS-treated conditions, we found five transcription factor genes showing decreased expression coupled with local chromatin compaction: Ptx1, caudal (cad), Senseless-2 (sens-2), odd and CG34376. All five were validated by RT-qPCR as downregulated upon trx knockdown (**Fig. 5D**) and may function as Trx-dependent repressors of EE fate regulators.

### Genetic validation of Trx-regulated transcription factors in repressing EE fate

To functionally validate transcription factors mediating Trx-dependent control of enteroendocrine (EE) differentiation, we first examined the expression of Trx-responsive candidates in the adult midgut using available GFP reporter lines and published data (Buchon et al., 2013; Dutta et al., 2015; Hang et al., 2020, Wu et al., 2021). All five factors were found to be expressed in intestinal cells, including stem cells, consistent with potential roles in lineage specification (Hung et al., 2020; **Suppl. Fig. 4A-B,D**)

RNAi-mediated depletion of candidate factors in esg⁺ progenitors revealed that sens-2, cad and Ptx1 tested lines caused elevated production of Pros⁺ EE cells compared with controls **(Suppl. Fig. 4C)**, with ptx1 knockdown eliciting the most pronounced increase. Furthermore, combined cad and trx knockdown produced a synergistic enhancement of the EE phenotype **(Suppl. Fig. 4C)**, supporting the idea that multiple Trx-responsive transcription factors cooperate to suppress EE fate, with ptx1 functioning as a central component of this network (**Figure 5E, Suppl. Fig 3C**).

Knockdown of *ptx1* in *esg⁺* progenitors led to a marked increase in Pros⁺ EE cells under both homeostatic (**Fig. 6A–B, C**) and DSS-induced regenerative conditions (**Fig. 6D–E, H**), effectively phenocopying *trx* RNAi. Conversely, overexpression of *ptx1* in progenitor cells suppressed EE differentiation and rescued the excess EE phenotype caused by *trx* depletion. Under DSS challenge, *ptx1* overexpression in *trx* RNAi intestines normalized Pros⁺ cell numbers and even prevented the hyperplastic accumulation of GFP⁺ progenitors typically observed following tissue damage (**Fig. 6F-H**). This result raises the possibility that Ptx1 interferes directly or indirectly with pathways that drive DSS-induced regenerative proliferation, such as Hippo signaling (Ren et al., 2010).

**Figure 6.**
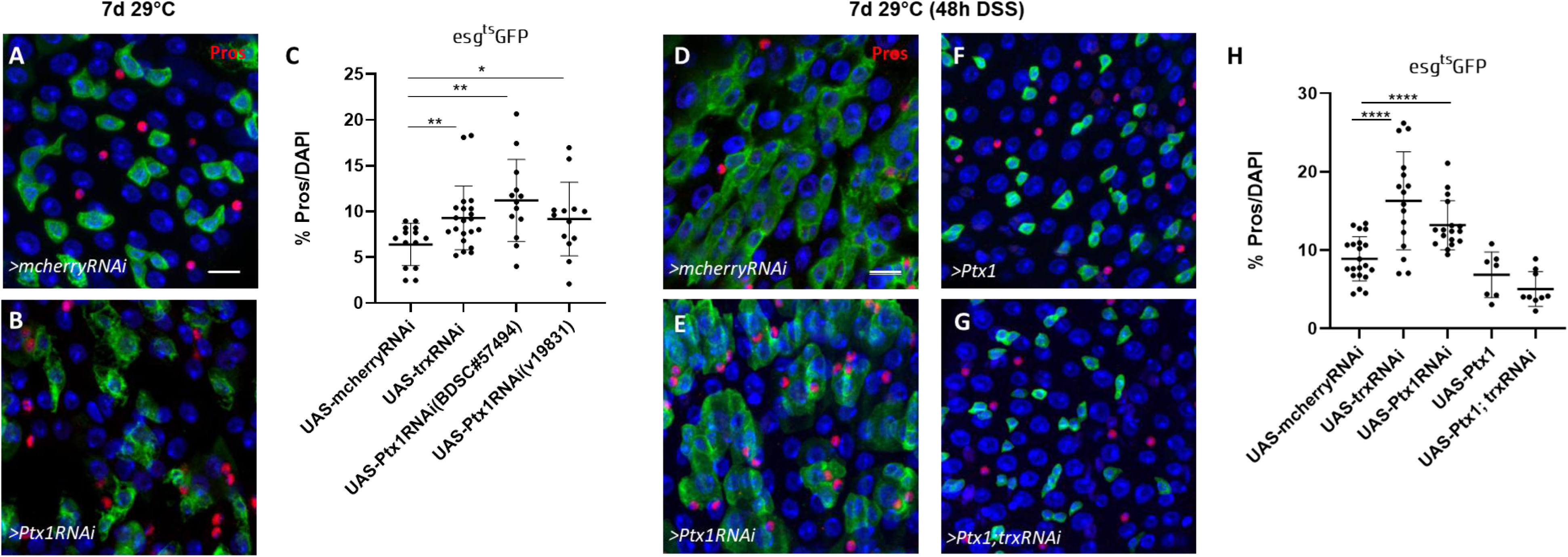
Genetic validation of Ptx1 as a Trx-dependent repressor of EE fate (A-B) Confocal images of esgts>GFP midguts crossed to control (mcherryRNAi) or UAS-Ptx1RNAi (BDSC#57494), showing a significant increase in the proportion of Pros+ enteroendocrine (EE) cells under homeostatic conditions 7 days upon RNAi induction at 29°C. Scale bars: 20 µm. (C) Quantification of Pros+ cells (% of DAPI-stained nuclei) for the genotypes shown in (A-B), using two independent Ptx1RNAi lines (BDSC#57494 and v19831) and UAS-trxRNAi (BDSC#33703). To generate the corresponding control genotype, mcherryRNAi was used as an additional control to ensure that the observed trxRNAi phenotype was not specific to the standard w1118 line. All measurements are representative of at least two independent experiments. Statistical analysis was performed using t-test; Data are shown as mean ± SD. (D-G) Immunofluorescence images from posterior midguts of young female flies carrying the esg-Gal4tsGFP driver and the indicated RNAi constructs. RNAi-mediated knockdown of trx and Ptx1 was performed using BDSC#33703 and BDSC#57494, respectively. Expression was induced at 29 °C for 7 days, with flies being exposed to DSS during the last 48 hours to enhance ISC activity. Single and double genotypes were generated from the same crosses using the identical RNAi lines. All genotypes were raised and handled under the same environmental conditions, and DSS treatment was applied uniformly before dissection. Enteroendocrine cells were identified by Prospero (red), with nuclei counterstained using DAPI (blue). All panels show a 20 µm scale bar. (H) Quantification of Pros⁺ cells (panels D–G) expressed as a percentage of total DAPI-stained cells. Data points correspond to individual zoom-in images. Statistical analysis was performed using a two-tailed unpaired t-test. Data are presented as mean ± SD.

Together, these data identify *Ptx1* as a key Trx-dependent transcriptional effector that is both necessary and sufficient to repress EE differentiation, under both homeostatic and regenerative conditions. We propose that Trx maintains progenitor identity by sustaining the expression of transcriptional repressors such as *Ptx1*, which constrain proneural gene activity and thereby prevent excess EE commitment.

## Discussion

Our study identifies Trithorax (Trx) as a chromatin regulator that safeguards ISC lineage fidelity by preventing inappropriate enteroendocrine (EE) differentiation. Trx achieves this by maintaining expression of transcription factors such as *Ptx1*, which restrain activation of the proneural transcriptional program and thereby stabilize progenitor identity to favour the EB-EC lineage. This adds a new dimension to the established paradigm of PcG/TrxG balance in intestinal homeostasis, where Polycomb complexes silence differentiation programs and other TrxG proteins (Kis, Trr) maintain gene activation required to restrict ISC proliferation (Veneti et al., 2024; Gervais et al., 2019).

### A reversible chromatin switch controlling ISC fate

A central question in stem cell biology is how a single stem cell type can generate distinct division outcomes over time. One possibility is chromatin remodeling through modulation of PRC/Trx activity, which would simultaneously affect multiple regulatory loci and initiate alternative developmental programs, such as EE versus EC fate. This is analogous to the progressive chromatin remodeling that occurs in neuroblasts, which is driven by temporal transcription factors and enables them to generate different types of neurons/ glia at different developmental stages. However, unlike neuroblast temporal transitions, which are irreversible (Holguera and Desplan, 2018), the ISC switch appears transient and reversible, allowing dynamic modulation of lineage output in response to physiological cues.

We propose that the EEP versus EB decision is exquisitely sensitive to Trx dosage. Even partial loss of Trx (*trx/+*) leads to excess EE cells (Tauc et al., 2021; **Fig. 2F**), implying that subtle shifts in the Trx/PRC equilibrium can alter cell fate. In the wild type, a modest upregulation of PRC2 activity could reduce Trx-dependent H3K4me1/2 and H3K27ac at key loci such as *Ptx1*, decreasing their expression. This would relieve repression of scute or the *phyllopod*–*Sina*–*Ttk* positive feedback circuit that amplifies its activity, enabling scute to cross the threshold required for an EEP-producing division. Subsequent restoration of Trx activity would reinstate *Ptx1* expression and return the ISC to its default EB-producing state. External stimuli such as mechanical stress or calcium influx into the cytoplasm (He et al., 2018) may transiently perturb this chromatin balance, triggering EE production during regeneration or aging.

### Trx-dependent enhancer regulation at the Ptx1 locus

The sensitivity of *Ptx1* to Trx dosage is consistent with genomic data showing that its enhancer–promoter region carries Trx-mediated active chromatin marks, including H3K4me2, which overlap with a Polycomb response element (Rickels et al., 2016). PRC1 (Pc) remains bound to this site whether the gene is active or repressed, while H3K4me2 and H3K27ac levels fluctuate in response to Trx activity. This arrangement—where enhancers and PREs are closely juxtaposed—renders *Ptx1* highly responsive to small chromatin shifts. Consistent with this view, our previous analysis of E(z)-dependent chromatin landscapes revealed an H3K27me3 peak at the *Ptx1* locus, supporting the idea that *Ptx1* is also subject to PRC2-mediated repression (Veneti et al., 2024). Thus, the *Ptx1* locus likely functions as a reversible chromatin switch that integrates Trx/PRC balance to determine whether an ISC produces an EB or an EE.

Published binding datasets (modENCODE Consortium et al., 2010) do not indicate direct *Ptx1* binding at the *scute* locus, but instead show association with *phyllopod*, a component of the *Phyl/Sina/Ttk* feedback loop that reinforces *scute* expression. With the caveat that these datasets do not come from intestinal progenitors, this could suggests that *Ptx1* restrains the EE program indirectly, by dampening this proneural positive feedback circuit rather than directly repressing *scute* itself.

### Coordination with other Trx-dependent transcription factors

Although *Ptx1* produced the strongest phenotype, RNAi against *sens-2*, *cad*, and *CG34376* also modestly increased EE numbers, indicating that multiple Trx-regulated transcription factors act in parallel to constrain EE differentiation. Combined *cad* and *trx* knockdown resulted in synergistic EE overproduction, supporting cooperative repression mechanisms *Cad* has been shown to inhibit JAK/STAT signaling, which promotes EC differentiation (Wu et al., 2021), suggesting it may act as a dual antagonist of both EC and EE fates.

Notably, four transcription factors identified in our genomic analysis (Ptx1, CG34376, cad and odd) exhibit regionalized expression along the anterior–posterior gut axis (Buchon et al., 2013; Hung et al., 2020) while maintaining lower basal expression throughout the epithelium. Such compartmentalized yet overlapping expression patterns resemble Hox gene domains, known targets of PcG/TrxG regulation, further highlighting their sensitivity to chromatin-dependent regulation. It is possible that these TFs collectively maintain EE fate repression along the entire length of the intestine.

### Concluding model

In summary, our data support a model in which Trx maintains ISC plasticity and lineage fidelity by sustaining the expression of *Ptx1* and other transcriptional repressors that counteract proneural activation. *Ptx1* carries Trx-mediated H3K4me2 marks at enhancer–PRE regions, making its expression exquisitely sensitive to the local Trx/PcG balance. When Trx levels drop, *Ptx1* expression falls, *phyllopod* and *scute* become derepressed, and ISCs undergo EE-producing divisions. Indeed, a small subset of ISCs (∼15%) has been reported to express higher levels of Sc at homeostasis (Chen 2018). Restoring Trx function reactivates *Ptx1* and silences the proneural circuit, returning ISCs to EB-producing mode (**Fig. 7**). This reversible chromatin switch provides a simple yet powerful mechanism by which adult stem cells can integrate epigenetic and physiological signals to flexibly alternate between differentiation programs while preserving tissue homeostasis.

**Figure 7.**
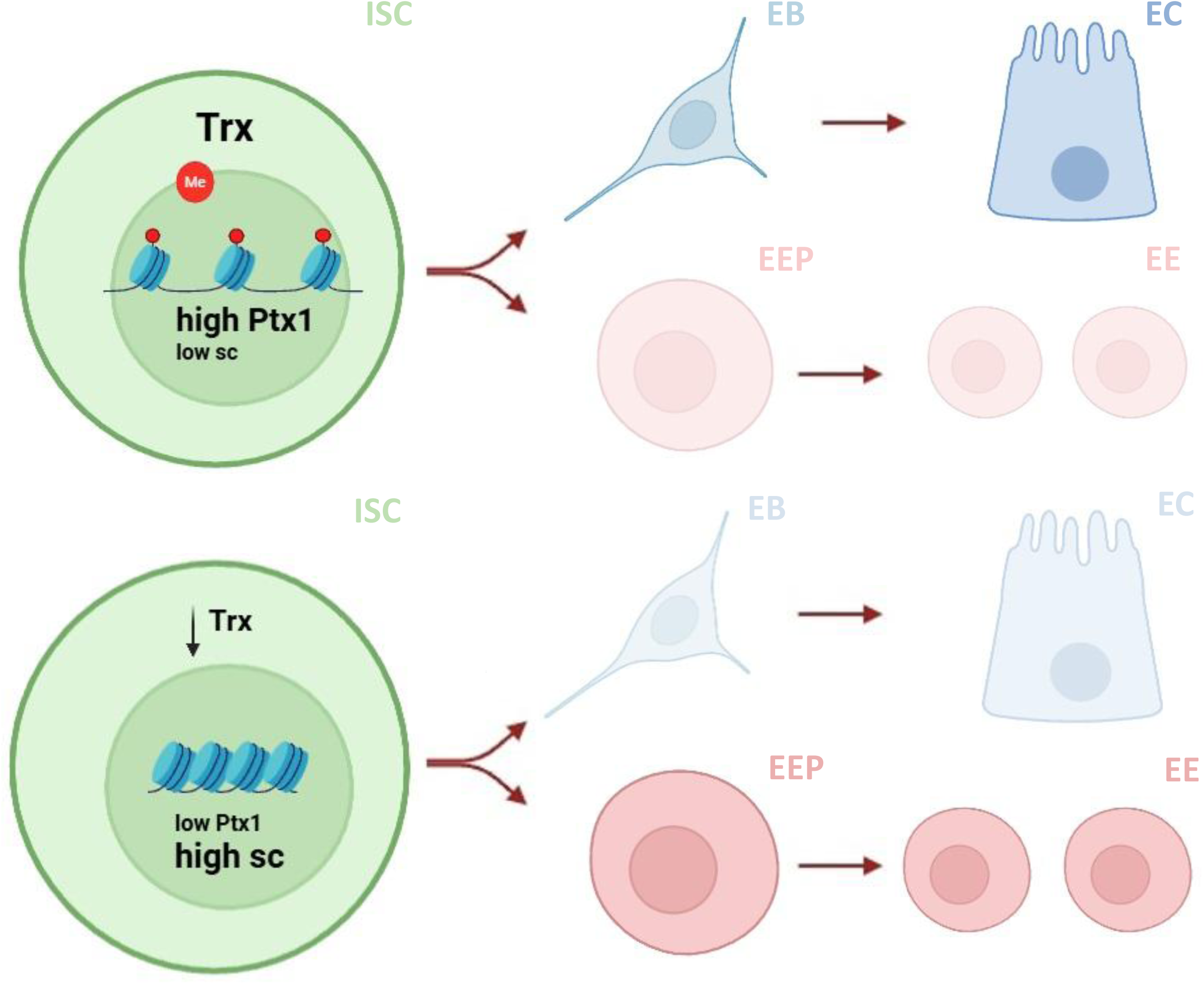
Proposed model Trx maintains ISC plasticity by sustaining Ptx1 expression, which opposes proneural activation. Loss of Trx reduces Ptx1 expression, leading to upregulation of scute and triggering EE-producing divisions. Restoring Trx reactivates Ptx1 and silences the proneural program, returning ISCs to EB-producing mode. This reversible chromatin switch enables adult ISCs to integrate epigenetic and physiological cues to balance differentiation and maintain tissue homeostasis.

Interestingly, MLL1, the mammalian Trx homolog, regulates *Pitx1 and Pitx2* in mouse intestinal stem cells (Goveas *et al*., 2021), mirroring the Trx-dependent activation of *Ptx1* we observe in flies. This conservation suggests a fundamental chromatin-transcription circuit in which Trx/MLL1 proteins stabilize Ptx1/PITX expression to maintain stem cell identity and fate.

## Acknowledgments

We thank the BDSC and VDRC for providing *Drosophila* stocks, the DSHB for antibodies, and FlyBase for genetic information. We also thank the IMBB Fly, Confocal, FACS-sorting and Genomics Facilities for their continuous support. This work was funded by the Hellenic Foundation for Research and Innovation (HFRI, Grant No. 1561) to Z.V. and Worldwide Cancer Research grant (WWCR, Grant No. 22-0345) to C. D.

## Contributions

V.F., A.G.E., C.D. and Z.V. conceived the project and designed the experiments. V.F. performed experiments and analyzed data. C.D. and Z.V. supervised the *Drosophila* genetic experiments and analyzed data. A.G. and P.H. performed and analyzed the RNA-seq and ATAC-seq experiments. V.T. performed and analyzed the ATAC-seq experiments. A.G.E. analyzed data and reviewed the manuscript. V.F., C.D. and Z.V. wrote the manuscript. All authors read and approved the final version.

## Materials and Methods

### Fly husbandry and genetics

All Drosophila genes used in this study are listed in FlyBase (http://flybase.bio.indiana.edu). Drosophila stocks and genetic crosses were maintained at 18 °C on standard cornmeal–yeast food to prevent premature activation of UAS-driven transgenes. Flies were kept under standard laboratory conditions with controlled light and humidity within the IMBB Insect Facility. Only mated female flies were used, as they possess larger midguts and exhibit distinct regenerative responses compared to males and virgin females, particularly after tissue damage.

To achieve temporally controlled gene expression, we employed the binary GAL4–GAL80^ts–UAS system. For overexpression and RNAi-mediated knockdown experiments, virgin female flies carrying the esg-Gal4, UAS-GFP, tubP-Gal80^ts, esg-Gal4^ts^ >GFP; Su(H)GBE-Gal80 or Su(H)GBE-Gal4^ts^ >GFP transgenes were crossed to either UAS or UAS-RNAi lines, or to appropriate control lines (w^1118 or UAS-mCherry-RNAi), as indicated in the figure legends. Crosses were maintained at 18 °C, and 3–4-day-old mated female F1 progeny were shifted to 29 °C for specific durations to induce transgene expression. The following RNAi and overexpression lines were obtained from the Bloomington Drosophila Stock Center (BDSC): UAS-mcherry-RNAi (BDSC#35785) UAS-trx-RNAi (BDSC#33703, #31092), trx^E2 (BDSC#24160), UAS-sc-RNAi (BDSC#26206), UAS-phyl-RNAi (BDSC#29433), UAS-ttk69 (BDSC#27325), UAS-Ptx1-RNAi (BDSC#57494), UAS-cad-RNAi (BDSC#57546, #34702), UAS-sens-2-RNAi (BDSC#27285, #34984), UAS-CG34376-RNAi (BDSC#29371), UAS-Notch (BDSC#5830). Additional fly lines were obtained from the Vienna Drosophila Resource Center (VDRC): w^1118 (v60000), UAS-trx-RNAi (v37715), UAS-Ptx1-RNAi (v19831). Unless otherwise stated, all genetic and genomic experiments involving trithorax (trx) knockdown were performed using the BDSC#33703 RNAi line.

For experiments employing the Su(H)-Gal4 driver, the UAS-trx-RNAi (v37715) line was used instead, as the BDSC#33703 stock led to pharate adult lethality even at 18 °C when combined with this driver.

Prior to dissections, adult flies were fed for 2 days with either 5% sucrose (S0389 Sigma-Aldrich) or 3% DSS (dextran sulfate sodium; MP Biomedicals) dissolved in 5% sucrose, delivered via soaked filter paper in empty vials, unless otherwise indicated. Progenitor cell populations were assessed by visualizing GFP-positive cells. GFP intensity was evaluated qualitatively; no quantitative fluorescence measurements were performed, as GAL4 titration effects in RNAi backgrounds could affect GFP levels.

To trace the lineage and assess the progeny of individual stem cells upon trx depletion, we employed MARCM (Mosaic Analysis with a Repressible Cell Marker) and G-TRACE systems. For MARCM clone generation, virgin females of genotype yw, hsFLP, tubP-Gal4, UAS-GFP/FM7; tubP-GAL80 FRT82B/TM6B were crossed either to w; FRT82B πMycx2 (control) or to y[1]; P{ry[+t7.2]=neoFRT}82B trx[E2]/TM6C (BDSC#24160) to generate trx mutant clones. Heat shock was performed for 1 hour at 37 °C on 3- to 4-day-old mated female F1 progeny. Flies were then maintained at 25 °C, and clones were analyzed 12 days post-induction.

G-TRACE lineage tracing was performed using esg-Gal4, tub-Gal80^ts to label progenitor cells with RFP and their immediate progeny with nuclear EGFP. This was achieved through expression of UAS-FLP, UAS-RedStinger, and Ubi-p63E^FRT-stop-FRT-StingerEGFP (BDSC#28280). These flies were crossed to either w^1118 (control) or UAS-trx-RNAi (BDSC#33703). Young mated females (3–5 days old) carrying the appropriate transgenes were shifted to 29 °C for 6 days to induce expression. One day prior to dissection, flies were fed with either 5% sucrose or 3% DSS as described above. Newly differentiated enterocytes (ECs) were identified as large, polyploid, DAPI-positive cells with nuclear EGFP and low or undetectable RFP signal. Newly generated enteroendocrine cells (EEs) were defined as Prospero-positive cells with nuclear EGFP and similarly reduced or absent RFP signal.

### Immunostaining and Image Analysis

Midguts were dissected in ice-cold 1× phosphate-buffered saline (PBS) and fixed in 4% formaldehyde for 20 minutes at room temperature. For Delta staining, an additional methanol fixation step was performed following formaldehyde fixation. Samples were washed three times for 10 minutes each in 1x PBS buffer, then incubated in blocking solution (PT with 0.5% BSA) for 30 minutes at room temperature. Primary antibodies, diluted in PBT, were applied overnight at 4 °C. The next day, samples were washed three times (10 min each) in PT buffer (1× PBS with 0.2% Triton X-100), incubated with secondary antibodies diluted in PBT for 1 hour at room temperature in the dark, and then washed again three times in PT before being mounted in Drop-n-Stain EverBrite™ Mounting Medium with DAPI (Biotium).

The following primary antibodies were used: mouse anti-Prospero (MR1A, 1:100, DSHB, RRID:AB 528440), rabbit anti-phospho-Histone H3 (Ser10), 1:1500, Merck Millipore, 06-570), mouse anti-Delta (C594.9B, 1:100, DSHB), rabbit anti-GFP (Minotech, Cat# 701-1, 1:10000) and goat anti-GFP (Origene AB0020-500). Secondary antibodies Alexa Fluor ® 488 Goat anti-Rabbit IgG (Thermo Fisher Scientific Cat# A-11034, RRID:AB_2576217), Alexa Fluor ® 555 Goat anti-Mouse IgG (Molecular Probes Cat# A-21424, RRID:AB_141780), Alexa Fluor ® 555 Donkey Anti-Rabbit IgG (Molecular Probes Cat# A-31572, RRID:AB_162543), Alexa Fluor ® 633 Goat Anti-Mouse IgG (Molecular Probes Cat# A-21052, RRID:AB_2535719) were used at 1:1000 dilution in PBT.

Confocal images were acquired using a Leica SP8 microscope with a 40× objective at the IMBB Confocal Facility. All images were taken from the posterior R4 region of the midgut. Image processing and analysis were performed using Leica LAS X software (version 3.7.2.22383).

Quantifications were performed using the CellCounter plugin in Fiji. Graphical representation, statistical analysis, and survival curves were generated using GraphPad Prism 8. Statistical significance was assessed using an unpaired two-tailed Student’s t-test.

### Midgut dissociation, cell sorting, and RNA extraction qRT-PCR

Midgut dissociation and FACS sorting were performed following a modified version of the protocol by Dutta et al. In brief, 120–150 midguts from esg-Gal4, UAS-GFP, tubP-GAL80^ts flies crossed to either w1118 or trx-RNAi (BDSC#33703) were dissected in filtered, sterilized, ice-cold DEPC-treated 1× PBS. Tissues were kept on ice throughout the dissections. The tissues were immediately subjected to enzymatic digestion in 1 mg/ml elastase (Sigma-Aldrich, cat. no. E0258) for 1 hour at 25 °C. During incubation, samples were subjected to gentle agitation and pipetted every 15 minutes to facilitate tissue dissociation. Following enzymatic digestion, intestinal cells were gently centrifuged at 400 × g for 20 minutes at 4°C, resuspended in 500 ul of ice-cold DEPC-treated PBS, passed through 40 μm cell strainers (BD Falcon) and GFP-positive progenitor cells were sorted in 200ul ice-cold DEPC-treated PBS using a FACSAria III cell sorter (BD Biosciences) at the IMBB FACS and Sorting Facility.

Sorted progenitor cells were immediately centrifuged at 400 × g for 20 minutes at 4°C, PBS was carefully removed and the cell pellet was resuspended in 700 μl QIAzol lysis buffer (Qiagen). Total RNA was isolated using the miRNeasy Micro Kit (Qiagen) according to the manufacturer’s instructions, diluted in 12ul of RNAse-free water and stored at −80°C.

### qRT-PCR

cDNA synthesis was carried out with SuperScript III Reverse Transcriptase (Invitrogen), as per the manufacturer’s guidelines using 500ng of total RNA extracted as described above. Quantitative PCR was performed using the KAPA SYBR Green Fast Master Mix (Roche) on a QuantStudio 1 Real-Time PCR system (Thermo Fisher Scientific). Gene expression levels were normalized to RpL32, and relative expression was calculated using the ΔΔCT method. Primer sequences (5’→ 3’) used for gene expression analysis were either retrieved from FlyPrimerBank or sourced from published literature reporting validated primers for the respective genes.

Ptx1 F: CGCCATGTGGACCAATCTTAC Ptx1 R: CGAACGGCTGCATGAACTG Sens-2 F: CTGCTTCACCAGCGCATACA Sens-2 R: TCGTCGGAGTTTCGCTTCTTG Cad F: AGCCGCCATACTTCGACTG

Cad R: TTATCCTTGGTGCGGGTTTTG Phyl F: CGTTCAGCTAATCCAGGCGAA Phyl R: GCCTCATTGCTGTTGACCG

Odd F: AGCAGAGAACAAGACTTTGCTG Odd R: TCCTCGCTTATCGCCGGAT

CG34376 F: CCCACAAGATAGTGCTGGCTA CG34376 R: CCACTCCTTGGTACATAAACTCC Ac F: TTTTCAACGACGACGAGGAG

Ac R: ACCATTGCTTAAATCGGCTA Sc F: AATGTAGACCAATCCCAGTCG Sc R: CACCACCCTTTGTCAAATCC

Lsc F: TCAAACTGTGGTAAACTCGCTGTC Lsc R: TCGGCGGAATTGTAGATGTG

Ase F: GCACAACCAGCAGAATCAAC

Ase R: AGGCAAACCCTTTCTTCCAG

Pros F: GCCCTGTTCCAACCACAATC

Pros R: CGAAACTGGTGAGTTGCTCG

Ttk F: ACAGCCTCCCAATGAAGATG

Ttk R: AGGTTGCTCTGGTGGTTGTT

Trx F: AATGCGGCGCGTTTCATTAA

Trx R: TGATGATGTGCTTGTGCCCT

Rpl-32 F: AGCATACAGGCCCAAGATCG

Rpl-32 R: GCACCAGGAACTTCTTGAATCC

### Sample preparation for genomics

For the genomic experiments, young virgin females carrying the esg-Gal4, UAS-GFP, tubP-Gal80^ts were crossed to either UAS-trxRNAi (BDSC#33703) or to w1118 control line.

150 young (3-4 days old) mated females from F1 progeny were shifted to 29 °C for 7 days to induce trx-RNAi expression. Two days prior dissection they were treated with either 5% sucrose or 3% DSS (dissolved in 5% sucrose) after which they were dissected in filtered ice-cold DEPC-treated 1× PBS and immediately dissociated with 1 mg/ml elastase (Sigma-Aldrich). Progenitor cells were sorted as GFP+ cells in a FACSAria III cell sorter, as indicated above. 50.000 GFP+ progenitors were sorted in ATAC buffer and approximately 200.000 GFP+ progenitors were sorted in QIAzol lysis buffer, following total RNA isolation using the miRNeasy Micro Kit (Qiagen).

### RNA-sequencing

RNA concentration was measured using a Qubit-4 Fluorometer (Invitrogen) with the Qubit RNA HS Assay Kit (Invitrogen) and RNA quality was assessed on an Agilent Tapestation 4150 with the High Sensitivity RNA ScreenTape (Agilent), according to manufacturers’ instructions. Libraries were prepared using the QuantSeq 3‘mRNA-Seq Library Prep Kit (Lexogen). Briefly, up to 150 ng of RNA were used for first strand synthesis, followed by RNA template removal and second strand synthesis initiated by random primers containing compatible linker sequences at their 5’ end. In-line barcodes were introduced during second strand synthesis, followed by magnetic bead-based purification. The resulting libraries were amplified for up to 19 cycles and purified. Library quantities were assessed using a Qubit 4 Fluorometer and the Qubit dsDNA HS Assay Kit (Invitrogen) and quality was assessed using the Agilent Tapestation 4150 and the High Sensitivity D1000 assay kit, according to manufacturers’ instructions. The quantified libraries were pooled equimolarly, and processed for Adapter Conversion PCR Amplification using the Universal Library Conversion Kit (App-A) (MGI Tech Co., Ltd.) according to manufacturer’s instructions. The quantity of the purified adapter conversion PCR (AC-PCR) product was assessed using a Qubit 4 Fluorometer and the Qubit dsDNA HS Assay Kit (Invitrogen) and the quality was evaluated using the Agilent Tapestation 4150 and the High Sensitivity D1000 assay kit, according to manufacturers’ instructions. This was followed by DNA denaturation, single-strand circularization, enzymatic digestion, enzymatic digestion product cleanup and quality control steps on the AC-PCR product, according manufacturer’s instructions. The resulting ssCirDNA was used for DNB preparation and sequencing on a DNBSEQ-G400 platform at the Genomics Facility of BSRC Alexander Fleming, using a G400 App-A FCS SE100 High-throughput Sequencing Set (MGI Tech Co., Ltd.).

FASTQ files were aligned against the drosophila reference genome build dm6 using a two-step approach. Briefly, reads were mapped to the reference transcriptome with HISAT2 and the remaining unmapped reads were aligned with bowtie2. Read counting on the 3’ UTRs was conducted using the metaseqR2 bioconductor package (Fanidis and Moulos, 2021). Read counting and filtering were conducted with default parameters, and normalization was carried out with DESEQ as implemented in metaseqR2.

For differential expression analysis, the PANDORA algorithm (Moulos and Hatzis, 2015) was used to estimate a meta p-value integrating the results from the DESEQ, DESEQ2, edgeR, limma, NBPSeq, and NOISeq algorithms. All additional parameters were set to default. Genes with meta p.value < 0.05 and absolute log2(FC) > 0.5 were considered as differentially expressed.

### ATAC- sequencing

ATAC-seq libraries were manufactured according to the original protocol (Buenrostro JB, et.al 2015). Briefly, 50-75K GFP positive sorted cells were collected in ATAC lysis buffer (10mM Tris-HCl ph 7.4, 10mM NaCl, 3mM MgCl2, 0.1% Igepal CA-630). Cells were pelleted 10’ x 500g, 4oC, supernatant discarded and subsequently gently suspended in 700ul ice-cold ATAC lysis buffer. After an additional 10’ x 500g, 4oC spin, lysis buffer was removed with a 1000p pipette leaving 22.5ul covering the nuclei. We added 2.5ul Tagmentation enzyme and 25ul 2xTagmentation Buffer (Illumina Tagment DNA enzyme and buffer kit (REF 20034210). Fragmentation reactions were performed using a bioSan TS-100 Thermo-Shaker at 37oC for 40’ shaking at 800RPM. Initial 5 cycle PCR was performed with Phusion High-Fidelity PCR Master Mix (NEB M0532S) using the Illumina Set C plate (REF 20026934) in total volume of 50ul. 5ul were used to estimate product cycle saturation with SYBR green in Biorad qPCR. Final libraries were produced after 13-14 total cycles of amplification and analyzed on a Bioanalyzer DNA chip. Libraries were sequenced on a Nextseq500, single-end 75bp reads, in the IMBB Genomics Facility.

Firstly, fastq files were quality trimmed with Trim Galore with the option “--max_n 5” to remove reads containing more than 5 ambiguous bases (https://github.com/FelixKrueger/TrimGalore).

Trimmed reads were aligned to the dm6 build with bowtie2 using the “--local --very-sensitive-local” parameters. Duplicates were discarded with Picard (http://broadinstitute.github.io/picard.).

Peak calling was performed with MACS3 using the parameters “--nomodel --shift −100 --extsize 200”. Differential accessibility analysis was conducted using the bioconductor package DiffBind. Briefly, peaks overlapping blacklist regions were removed, and only peaks present in at least two samples with more than 10 counts were retained. Read extension and summit re-centering were disabled. Differentially accessible regions were identified using DESEQ2 with an absolute fold change greater than 0.5 and FDR < 0.05. Peak-to-gene annotation was performed with bedtools (https://bedtools.readthedocs.io/en/latest/), assigning each peak to genes with transcription start sites (TSSs) located within ±10 kb of the peak center or overlapping the peak body.

---

**Supplementary Figure 1.**
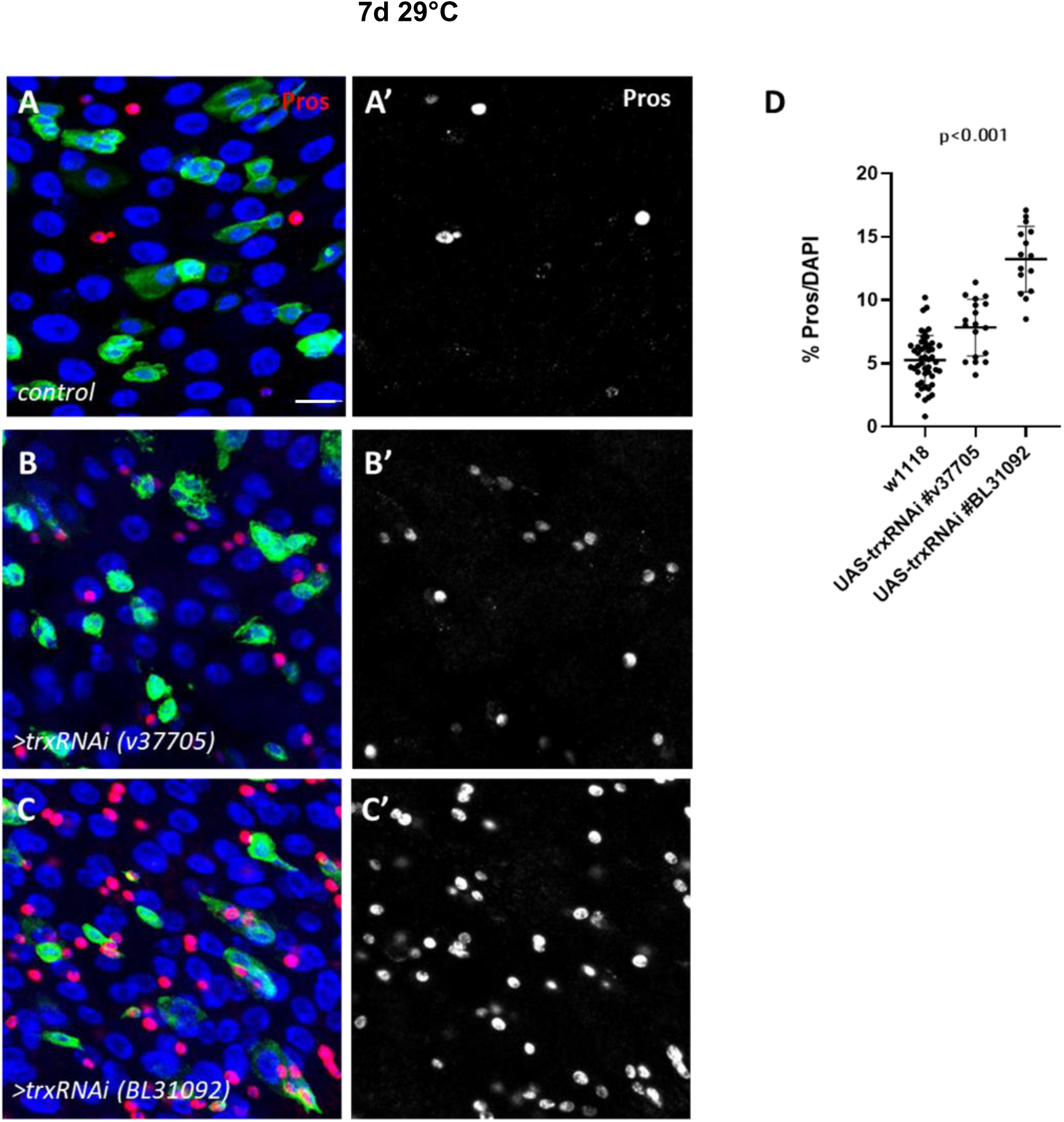
(A–C) Confocal images of midguts from flies expressing esg^ts>GFP crossed with two independent trxRNAi lines (v37715 and BDSC#31092) or control (w1118), showing the increased number of Pros+ enteroendocrine (EE) cells upon trx knockdown. (D) Quantification of Pros+ cells as a percentage of total DAPI-stained nuclei from the genotypes shown in (A–C). Each point represents an individual zoom-in image; Data indicate mean ± SD.

**Supplementary Figure 2.**
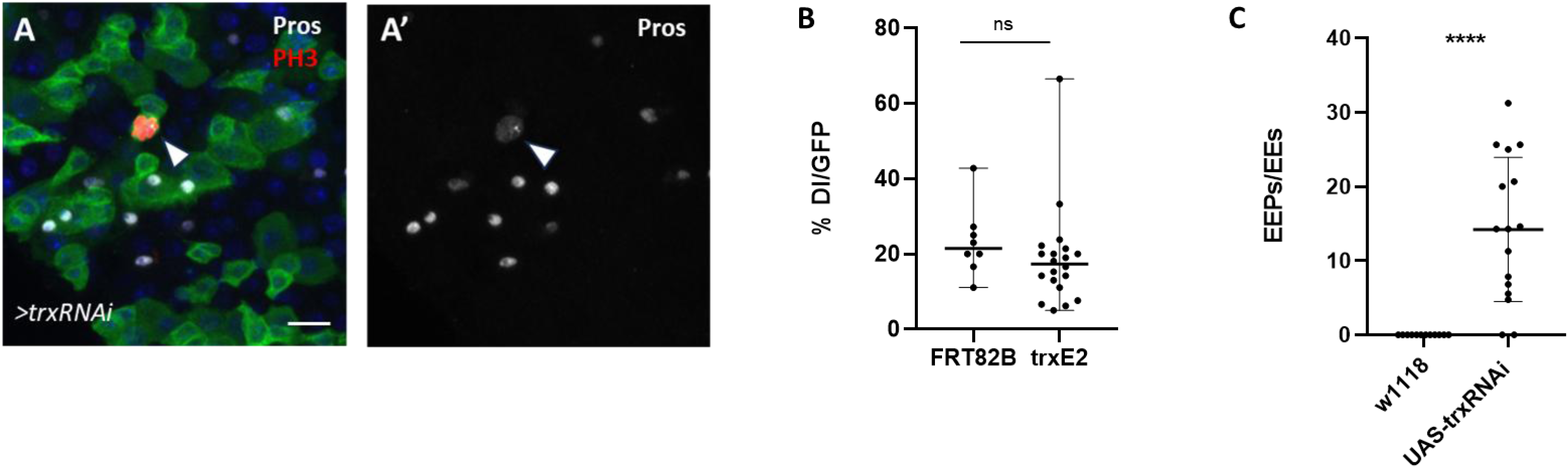
(A–A’) Confocal images of esgts>trxRNAi midguts (BDSC#33703) stained for PH3 (mitotic cells, red) and Pros (EE marker, white). Arrows indicate PH3+ cells that also stain for Pros, consistent with enteroendocrine precursors (EEPs). (B) Quantification of Dl-positive intestinal stem cells (ISCs) as a percentage of GFP+ cells from MARCM clones carrying the trx^E2 amorphic allele, showing that ISC numbers are unchanged upon trx depletion. (C) Quantification of newly generated EE progenitors (EEPs, Pros⁺GFP⁺ cells) as a fraction of total Pros⁺ EEs, reflecting recent ISC divisions biased toward the EE fate. The graph corresponds to the immunostainings shown in Figure 3, panels M and N, where aged control and trxRNAi flies (BDSC#33703) were subjected to DSS-induced epithelial damage.

**Supplementary Figure 3.**
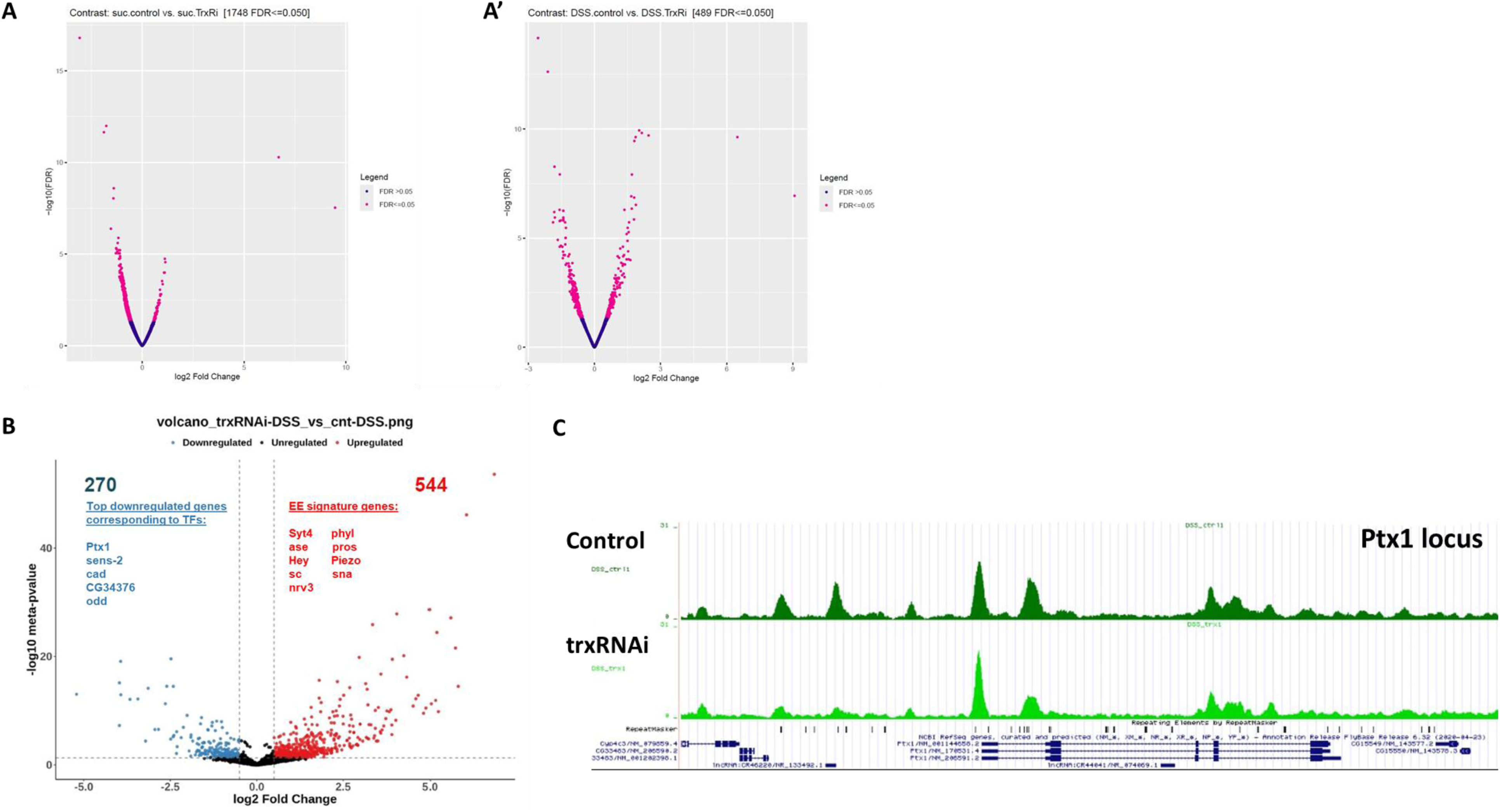
(A-A’) Volcano plot of ATAC-seq data comparing control and trxRNAi FACS-sorted progenitors under homeostasis (A) and DSS treatment (A’). Peaks with nominal significance (FDR ≤ 0.05) are shown in pink. A total of 1,748 and 489 chromatin regions were significantly differentially accessible in sucrose and DSS conditions, respectively. (B) Volcano plot of RNA-seq data depicting genes significantly up- or down-regulated upon trx knockdown in FACS-sorted progenitors under DSS-induced epithelial stress. Up-regulated transcripts are shown in red, down-regulated in blue. A subset of up-regulated genes associated with EE fate is highlighted in red, and a subset of down-regulated genes corresponding to TFs is highlighted in blue. (C) UCSC Genome Browser view of the Ptx1 locus showing normalized ATAC-seq signal in control and trxRNAi ISC progenitors under DSS feeding. Ptx1 is among the loci commonly exhibiting reduced chromatin accessibility in trxRNAi progenitors under both sucrose and DSS conditions, with decreased accessibility extending into upstream regulatory regions.

**Supplementary Figure 4.**
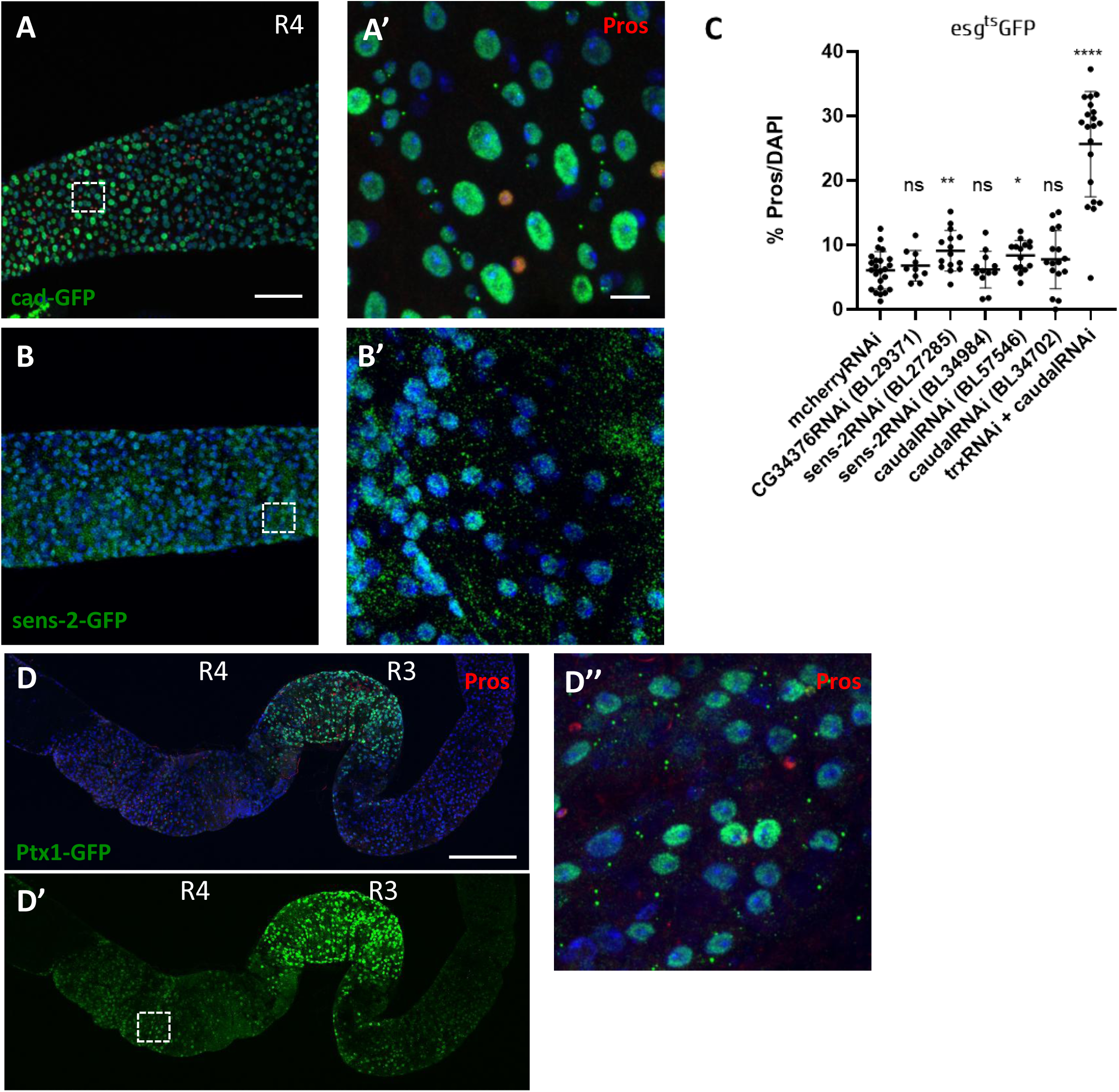
(A-A’) Expression of the cad-GFP reporter reveals cad expression throughout the R4 region of the intestinal epithelium, detected in all intestinal cell types. Prospero marks enteroendocrine (EE) cells, and nuclei are counterstained with DAPI. Scale bars: 50 µm (overview) and 20 µm (zoom-in). (B-B’) Expression of the sens-2 reporter line is observed throughout the R4 region of the intestinal epithelium, encompassing all intestinal cell types, including small diploid cells. Nuclei were visualized with DAPI. Scale bars: 50 µm (overview) and 20 µm (zoom-in). (C) Quantification of Pros⁺ cells (expressed as percentage of total DAPI-stained cells) in midguts after RNAi knockdown of candidate EE fate repressors (CG34376, sens-2, and caudal), compared to control (UAS-mcherryRNAi). For the double knockdown of caudal and trithorax, the RNAi lines BDSC#33703 (trithorax) and BDSC#57546 (caudal) were used. This analysis tests whether depletion of the candidates in esg+ cells produces a Pros⁺ cell expansion reminiscent of the phenotype observed in trx-depleted progenitors. Statistical comparisons between each RNAi line and the control are shown above the corresponding bars. Data are shown as mean ± SD; significance was determined by a two-tailed unpaired t-test. (D-D’’) Expression pattern of the Ptx1-GFP reporter line indicates strong expression of Ptx1 in the R3 region of the midgut epithelium and a weaker but still detectable expression in the R4 region of the posterior midgut. Prospero has been used as an EE cell marker and nuclei have been counterstained with DAPI. Representative area of middle and posterior midgut is shown. Scale bar; 150μΜ (Α-A’) and 20μΜ (Α’’).

